# Bone marrow mesenchymal stromal cells mediate cellular inflammation in HFpEF

**DOI:** 10.1101/2025.03.28.645924

**Authors:** Kyle I. Mentkowski, I-Hsiu Lee, David Rohde, Alexandre Paccalet, Hana Seung, Nina Kumowski, Maximilian J. Schloss, Michael Carpenito-Kronenfeld, Vaishali Kaushal, Ritika Jain, Xhoana Memcaj, Jacob Wiebe, Noor Momin, Steffen Pabel, Charlotte G. Muse, Kaysaw Tuy, Sarah Ung, Jason Roh, Filip K. Swirski, David Scadden, Peter van Galen, Antonia Kreso, Kamila Naxerova, Maarten Hulsmans, Matthias Nahrendorf

## Abstract

During the genesis of heart failure, the myocardium recruits an abundance of bone marrow-derived leukocytes, primarily monocytes, with various disease-promoting functions. Increased hematopoiesis fuels these unfavorable changes in cardiac leukocyte origin, number and phenotype. Here we examine hematopoietic niche cells, which regulate blood progenitor proliferation and systemic monocyte supply, in obese, hypertensive mice that develop heart failure with preserved ejection fraction (HFpEF). Single cell transcriptomics revealed that in HFpEF, stromal bone marrow niche cells expand and respond strongly to IFNɣ. Deleting the IFNɣ receptor in stromal cells of *Prrx1*^CreERT2^;*Ifngr1*^fl/fl^ mice reduced hematopoietic progenitor proliferation and systemic monocytes in both the steady state and HFpEF and also increased the canonical hematopoietic maintenance factor CXCL12, resulting in reduced fibrosis and improved diastolic function. CD8+ T cells in adipose tissue were a major source of IFNɣ in mice with HFpEF; their depletion restored CXCL12 expression and lowered monocyte numbers. ScRNA-seq in mice with ischemic heart disease uncovered a diverging marrow response. These data indicate that in HFpEF, adipose tissue, bone marrow and adaptive and innate immune cells conspire to expand harmful macrophage subsets in the heart.

## Introduction

Heart failure with preserved ejection fraction (HFpEF) is an expanding burden to healthcare systems worldwide, accounting for more than half of all heart failure cases and affecting over 30 million patients.^1–3^ Morbidity and mortality rates for patients with HFpEF remain high.^4^ Prevalence of HFpEF is rising in association with an aging population and the continued expansions of obesity, diabetes, and hypertension.^5–7^ These comorbidities induce systemic inflammation and play major roles in the pathogenesis of HFpEF. Recent trials utilizing SGLT2 inhibitors ^8, 9^ have shown promise for treating HFpEF; however, there is still an urgent need for additional therapeutic options.

Emerging data indicate that the inflammation observed in individuals with obesity and hypertension is triggered in part by systemically expanding leukocytes and elevated hematopoiesis.^10–13^ A rise in circulating, pro-inflammatory leukocytes has been linked to increased cardiovascular events and mortality in HFpEF.^14^ These short-lived immune cells, particularly CCR2^+^ monocytes, can migrate into the heart in acute or chronic injury and influence the pathogenesis of cardiovascular disease.^15–19^ Hematopoietic stem and progenitor cells (HSPC) in the bone marrow supply recruited inflammatory cells.^20^ HSPC proliferation and the lineage of produced cells are principally regulated by niche factors in the bone marrow microenvironment.^21^ In response to infection, diabetes or cardiovascular disease, circulating factors can influence stromal cells in the bone marrow niche to direct HSC toward myeloid lineage bias.^22–25^

There is now substantial evidence that recruited macrophages contribute to inflammation and fibrosis in many forms of cardiovascular disease,^17, 26–28^ but the cellular and molecular pathways that amplify myelopoiesis in HFpEF induced by obesity and hypertension have not been studied systematically.^10^ Understanding the mechanisms behind increased leukocyte supply to the myocardium may provide avenues to dampen harmful inflammatory pathways in HFpEF. We therefore systematically examined the stromal bone marrow compartment in an established mouse model of cardiometabolic HFpEF caused by its most common risk factors, namely obesity and hypertension.^29^

## Results

### Increased monocytosis contributes to obesity- and hypertension-induced HFpEF

We began by studying changes to blood cell production and the myocardial immune cell repertoire in an established model of HFpEF elicited by high-fat diet (HFD) and L-NAME-induced hypertension.^29^ Assaying hematopoiesis by flow cytometry, we found higher numbers and fractions of actively cycling multipotent progenitor cells (MPP, Figure 1A-D), which contain monocyte-biased progenitors.^30^ No significant changes in the number and fraction of actively cycling CD48-CD150+ HSC (SLAM-HSC) were noticed (Figure 1A-D). In mice with HFpEF, monocytes increased in the bone marrow and in circulation (Figure 1E-H), thereby expanding Ly6C^high^ monocyte and macrophage abundance in the heart (Figure 1I and 1J). As expected,^29^ doppler echocardiography in HFpEF mice displayed multiple indicators of diastolic dysfunction with preservation of left ventricular ejection fraction and no change in heart rate when compared with age-matched control mice (Figure 1K and 1L). In line with elevated HSPC cycling, bone marrow plasma levels of CXCL12, a key hematopoietic quiescence factor supplied by mesenchymal stromal cells (MSC),^31, 32^ were decreased in HFpEF mice (Figure 1M). In the sternal bone marrow of HFpEF patients, a similar decrease in CXCL12 levels was found (Figure 1N, see Table S1 for clinical characteristics). These results highlight a hematopoietic niche-driven rise in both myelopoiesis and monocytosis which occur in parallel with worsening diastolic function in obese mice with hypertension.

**Figure 1:**
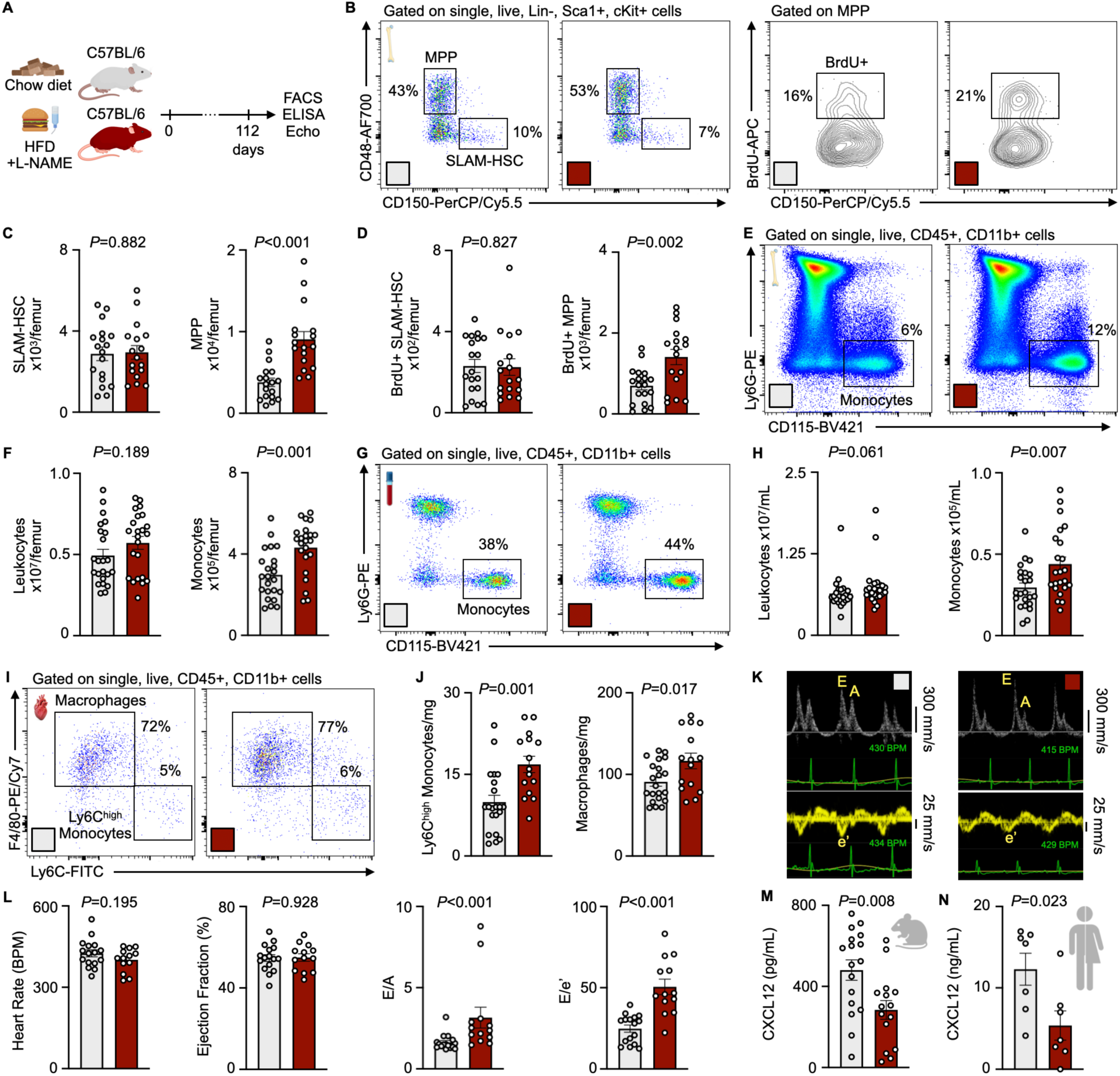
Increased myelopoiesis and monocytosis in HFpEF. (**A**) Experimental outline of HFpEF mouse model. (**B**) Gating strategy to identify SLAM-HSCs and MPPs, as well as the percentage of each population that is proliferating (BrdU+). (**C-D**) Number of total and proliferating SLAM-HSCs and MPPs in the femora of control and HFpEF mice after 16 weeks of HFD (high fat diet) and L-NAME, as measured by flow cytometry (n=17-19 mice per group, unpaired student’s t-test). (**E**) Gating strategy to identify bone marrow monocytes. (**F**) Numbers of leukocytes and monocytes in the bone marrow of control and HFpEF mice (n=22-23 mice per group, unpaired student’s t-test or Mann-Whitney test depending on normality). (**G**) Gating strategy for blood monocytes. (**H**) Enumeration of leukocytes and monocytes in the blood of control and HFpEF mice (n=22-23 mice per group, unpaired student’s t-test or Mann-Whitney test depending on normality). (**I**) Gating strategy to identify cardiac myeloid cells. (**J**) Cardiac macrophage and Ly6C^high^ monocyte numbers in the heart during HFpEF (n=19-22 mice per group, unpaired student’s t-test). (**K**) Representative pulsed-wave Doppler (upper) and tissue Doppler (lower) echocardiography tracings of control and HFpEF mice. (**L**) Quantification of echocardiography measurements (n=13-17 per group, unpaired student’s t-test or Mann-Whitney test depending on normality). (**M**) CXCL12 levels in bone marrow plasma of control and HFpEF mice (n=15-19 per group, unpaired student’s t-test). (**N**) CXCL12 levels in sternal bone marrow plasma of control and HFpEF patients (n=7 per group, unpaired student’s t-test).

Using genetic fate mapping in *Cx3cr1*^CreER^;*Ai9*^fl/fl^ mice (Figure S1), where resident macrophages are labelled as Td-tomato positive at the time point of our analysis, we found that Td-negative recruited cardiac macrophages were significantly increased in HFpEF compared with chow diet normotensive controls (Figure 2A and 2B). These findings support that in HFpEF, cardiac macrophage numbers expand due to recruitment of bone marrow-derived monocytes. We next compared cardiac leukocyte numbers and heart function in *Ccr2*^-/-^ mice versus C57BL/6 control mice with HFpEF (Figure 2C). *Ccr2*^-/-^ mice, in which monocyte migration is abrogated,^33, 34^ had significantly lower Ly6C^high^ monocyte and macrophage numbers in the heart, thus confirming that recruited monocytes contribute to these cell populations (Figure 2D). Reduction in recruited macrophages by *Ccr2* deletion improved diastolic function (Figure 2E). These data document that recruited macrophages contribute to diastolic dysfunction in HFpEF induced by obesity and hypertension.

**Figure 2:**
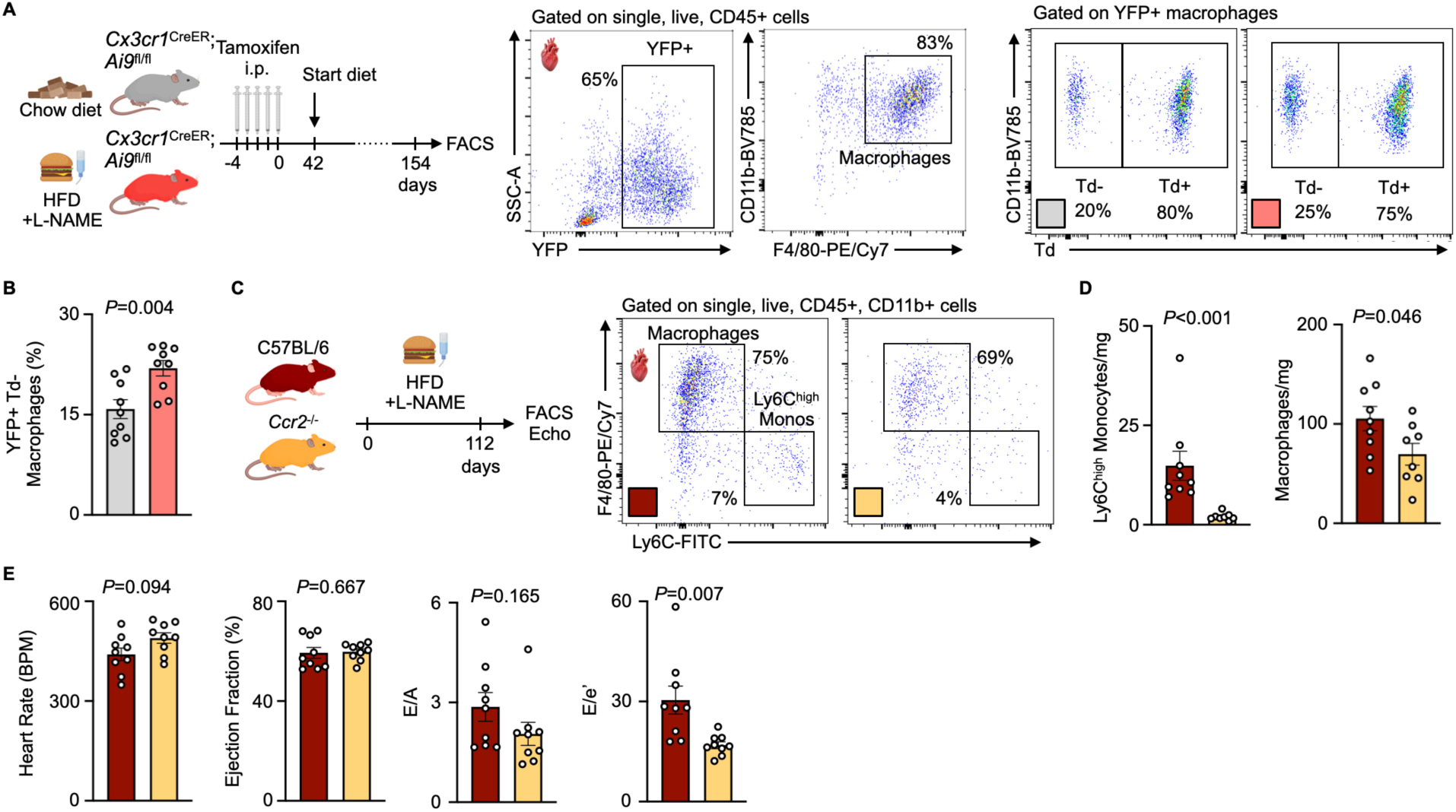
Recruited macrophages expand and contribute to HFpEF. (**A**) Experimental outline and gating strategy to identify YFP+ Td-cardiac macrophages in *Cx3cr1*^CreER^;*Ai9*^fl/fl^ mice in HFpEF vs. control settings. (**B**) YFP+ Td-cardiac macrophage frequency in control vs. HFpEF *Cx3cr1*^CreER^;*Ai9*^fl/fl^ mice (n=9 per group, unpaired student’s t-test). (**C**) Experimental outline of HFpEF in C57BL/6 vs. *Ccr2*^-/-^ mice. (**D**) Quantification of macrophage and Ly6C^high^ monocyte numbers in *Ccr2*^-/-^ HFpEF mice compared with C57BL/6 HFpEF controls (n=8-9 per group, unpaired student’s t-test or Mann-Whitney test depending on normality). (**E**) Parameters of diastolic function as measured by Doppler echocardiography in *Ccr2*^-/-^ HFpEF mice compared with C57BL/6 HFpEF controls (n=9 per group, unpaired student’s t-test or Mann-Whitney test depending on normality).

### Single cell transcriptomic profiling of the hematopoietic niche in HFpEF mice

Myeloid cell production is a tightly controlled process, orchestrated by hematopoietic progenitors and influenced by bone marrow stromal cells.^21^ To determine the pathways which increase pro-inflammatory myeloid cell production in HFpEF, we relied on unbiased single cell RNA sequencing (scRNA-seq) to profile bone marrow niche cells from HFpEF mice compared with those from control mice that were normotensive and consumed a chow diet (Figure 3A). After removing hematopoietic cells from the dataset, a total of 4,919 niche cells were obtained. Using known lineage markers, niche cells were divided into 13 niche cell clusters, including mesenchymal stromal cells (MSC), endothelial cells (EC), osteolineage cells (OLC), fibroblasts and pericytes (Figure 3B, Figure S2A-S2C).^35, 36^ Two major MSC clusters, Leptin-receptor+ MSC (LepR-MSC) and MSC 1, significantly expanded in HFpEF, while transitional EC were reduced (Figure 3C, Figure S2D). We confirmed this observation by flow cytometry, which underscored a higher number of MSC in HFpEF mice compared with controls (Figure 3D).

**Figure 3:**
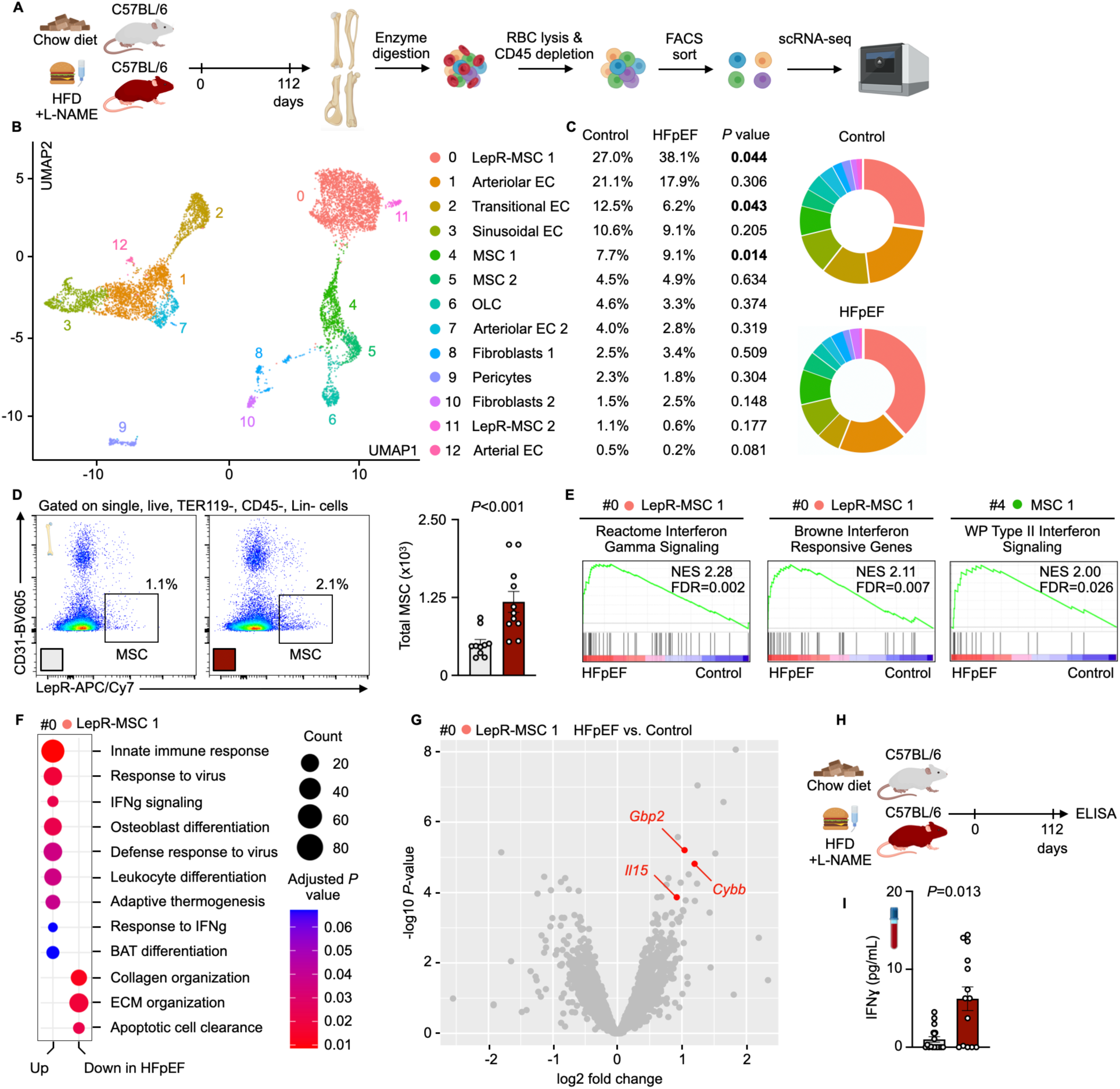
Single cell transcriptomics of the hematopoietic niche in HFpEF. (**A**) Experimental outline illustrating scRNA-seq analysis of bone marrow stromal cells from 8 control mice and 8 HFpEF mice, each pooled into 4 final replicates per group. (**B**) Dimensionality reduction analysis by uniform manifold approximation and projection (UMAP) shows 13 bone marrow stromal cell populations. (**C**) Relative cell population frequencies in control and HFpEF mice. Statistically significant changes in distribution determined by two-tailed Student’s t-test. (**D**) Flow cytometry gating for mesenchymal stromal cells. Bar graph shows increased MSC in HFpEF mice compared with control mice. Each data point represents the pooled bones of 1 mouse (n=10-11 per group, two-tailed Mann-Whitney test). (**E**) Gene set enrichment analysis (GSEA) plots highlighting interferon gamma signaling in LepR-MSCs (cluster 0) and MSC 1 (cluster 4) in HFpEF mice compared with control mice. (**F**) Significant gene ontology biological process (GOBP) gene sets, highlighting major up- or down-regulated gene sets in LepR-MSC (cluster 0) in HFpEF mice compared with control mice. Dot color indicates statistical significance and dot size the number of enriched genes. (**G**) Volcano plot of significantly altered genes in LepR-MSC (cluster 0). Marked genes are associated with interferon gamma signaling and have an FDR<0.05. (**H**) Experimental outline of serum collection from HFpEF vs. control mice. (**I**) Serum levels of IFNɣ protein, measured by ELISA, in HFpEF mice compared with control mice(n=15-20 mice per group, two-tailed Mann-Whitney test).

Because mice with HFpEF had expanded MSC, and based on their established involvement in maintaining hematopoietic stem cell function^37, 38^, we explored the transcriptional profile of MSC in greater detail. We identified 44 differentially expressed genes (DEGs) between HFpEF and control mice, using *FDR*<0.05 as the significance cutoff (Figure S2E-S2G). Differentially regulated genes were further analyzed by gene set enrichment analyses (GSEA), which revealed an interferon gamma (IFNɣ) signaling response in MSC (Figure 3E). Gene ontology biological process (GOBP) gene sets similarly highlighted a response to IFNɣ, as well as leukocyte differentiation pathways in MSC (Figure 3F). Of the top DEGs in LepR-MSC, IFNɣ response genes were among the most up-regulated genes (Figure 3G). We then analyzed blood serum by ELISA (Figure 3H) and found circulating IFNɣ protein was significantly elevated in HFpEF mice compared with controls (Figure 3I), a result that is consistent with augmented serum IFNɣ levels in a subset of HFpEF patients.^39^ Taken together, these data suggested a MSC-driven bone marrow response to IFNɣ in mice with HFpEF.

### IFNɣ sensing by MSC regulates steady-state hematopoiesis

Having established increased IFNɣ in the blood of mice with HFpEF as well as IFNɣ-activated pathways in their bone marrow niche cells, we wondered if IFNɣ-sensing by MSC can influence hematopoiesis. Of note, prior work indicated that IFNɣ directly stimulates HSPC proliferation^40, 41^ and can influence the secretome of human bone marrow MSC *in vitro*.^42^ Additionally, mice with IFNɣ overproduction demonstrated a myeloid-biased rise in hematopoiesis,^42^ which could be due to MSC-specific signaling or direct HSPC activation. We thus next generated *Prrx1*^CreERT2^;*Ifngr1*^fl/fl^ mice, which lack IFNɣ receptor expression in MSC, and first characterized these mice in steady state (Figure 4A). Successful gene deletion in MSC was confirmed by flow cytometry staining for the IFNɣ receptor (Figure S3A-C). In *Prrx1*^CreERT2^;*Ifngr1*^fl/fl^ mice, we detected no change in MSC number (Figure S3D) or blood leukocyte levels prior to tamoxifen administration, compared with *Prrx1*^CreERT2^ controls (Figure S3E and S3F). *Prrx1*^CreERT2^;*Ifngr1*^fl/fl^ mice had similar heart function (Figure S4) as *Prrx1*^CreERT2^ controls at baseline. Interestingly, both male and female *Prrx1*^CreERT2^;*Ifngr1*^fl/fl^ mice exhibited a significant decrease in both number and fraction of actively cycling SLAM-HSC (Figure 4B and 4C), resulting in a numerical decline of leukocytes and monocytes in the bone marrow (Figure 4D and 4E) and blood (Figure 4F). Using a colony forming unit (CFU) assay as an orthogonal approach (Figure 4G), we found that *Prrx1*^CreERT2^;*Ifngr1*^fl/fl^ mice formed fewer granulocyte/macrophage progenitor colonies (CFU-GM) in blood and bone marrow when compared with *Prrx1*^CreERT2^ controls (Figure 4H). Consistent with curtailed hematopoiesis, *Cxcl12* gene expression rose in FACS-sorted MSC and CXCL12 protein in bone marrow plasma from *Prrx1*^CreERT2^;*Ifngr1*^fl/fl^ mice compared with *Prrx1*^CreERT2^ controls (Figure 4I and 4J). Considered collectively, these data support that bone marrow MSC play a role in IFNɣ sensing and that IFNɣ serves as a tonic regulator of steady-state hematopoiesis.

**Figure 4:**
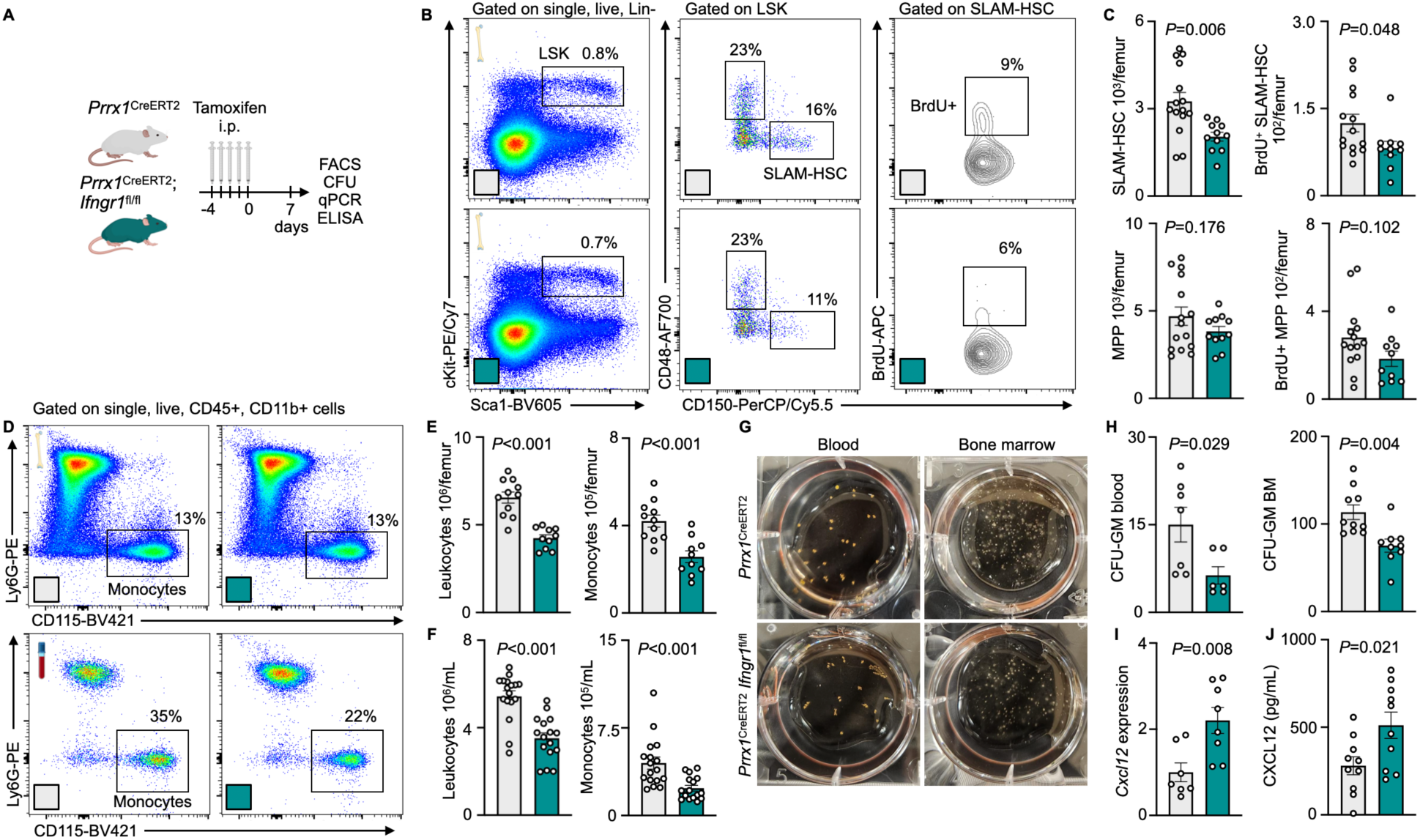
IFNɣ sensing by MSC regulates steady-state hematopoiesis. (**A-B**) Experimental outline and gating strategy to analyze hematopoiesis in *Prrx1*^CreERT2^ control vs. *Prrx1*^CreERT2^;*Ifngr1*^fl/fl^ mice in the steady state. (**C**) SLAM-HSC numbers and their BrdU incorporation are significantly reduced in *Prrx1*^CreERT2^;*Ifngr1*^fl/fl^ compared with *Prrx1*^CreERT2^ controls (n=11-16 per group, unpaired student’s t-test). (**D-F**) Flow cytometry gating and leukocyte numbers in the bone marrow and blood of *Prrx1*^CreERT2^;*Ifngr1*^fl/fl^ compared with *Prrx1*^CreERT2^ controls (n=10-17 per group, unpaired student’s t-test). (**G-H**) Images and colony enumeration of CFU assay in blood and bone marrow in *Prrx1*^CreERT2^;*Ifngr1*^fl/fl^ compared with *Prrx1*^CreERT2^ controls (n=6-10 per group, two-tailed Mann-Whitney test). (**I**) Gene expression in MSC sorted from *Prrx1*^CreERT2^;*Ifngr1*^fl/fl^ vs. *Prrx1*^CreERT2^ controls (n=7-8 per group, two-tailed Mann-Whitney test). (**J**) CXCL12 protein abundance in bone marrow plasma from *Prrx1*^CreERT2^;*Ifngr1*^fl/fl^ vs. *Prrx1*^CreERT2^ controls (n=10 per group, unpaired student’s t-test).

### Inhibiting MSC-specific IFNɣ sensing reduces monocytosis and improves cardiac function in mice with HFpEF

Next, we assessed hematopoiesis and cardiac function in *Prrx1*^CreERT2^;*Ifngr1*^fl/fl^ and *Prrx1*^CreERT2^ mice with HFpEF (Figure 5A). IFNɣ levels in circulation were unchanged between groups (Figure 5B). Consistent with steady-state measurements in *Prrx1*^CreERT2^;*Ifngr1*^fl/fl^ and *Prrx1*^CreERT2^ mice (Figure 4I and 4J), *Cxcl12* gene expression in FACS-sorted MSC and CXCL12 protein expanded in the bone marrow of *Prrx1*^CreERT2^;*Ifngr1*^fl/fl^ mice compared to *Prrx1*^CreERT2^ mice (Figure 5C and 5D). *Prrx1*^CreERT2^;*Ifngr1*^fl/fl^ mice with HFpEF had fewer HSC (Figure 5E and 5F) and monocytes in the bone marrow (Figure 5G and 5H) and blood (Figure 5I and 5J). In the heart, *Prrx1*^CreERT2^;*Ifngr1*^fl/fl^ mice exhibited similar numbers of total cardiac macrophages (Figure S5A and S5B) but smaller numbers of Ly6C^high^ monocytes and proportions of CCR2+ macrophages (Figure 5K-5M), thus demonstrating a shift toward fewer recruited macrophages. As a result, there were more resident macrophages in *Prrx1*^CreERT2^;*Ifngr1*^fl/fl^ mice with HFpEF compared to *Prrx1*^CreERT2^ mice with HFpEF (Figure S5A-S5C). Lessened macrophage recruitment was linked to decreased interstitial myocardial fibrosis (Figure 5N and 5O) and improved diastolic function (Figure 5P) in *Prrx1*^CreERT2^;*Ifngr1*^fl/fl^ mice with HFpEF compared to *Prrx1*^CreERT2^ mice with HFpEF. Of note, gene deletion in *Prrx1*^CreERT2^;*Ifngr1*^fl/fl^ mice did not affect cardiac fibroblast expression of the IFNɣ receptor (Figure S5D and S5E), thus supporting that the attenuated myocardial fibrosis and improved diastolic function in *Prrx1*^CreERT2^;*Ifngr1*^fl/fl^ mice resulted from reduced myelopoiesis.

**Figure 5:**
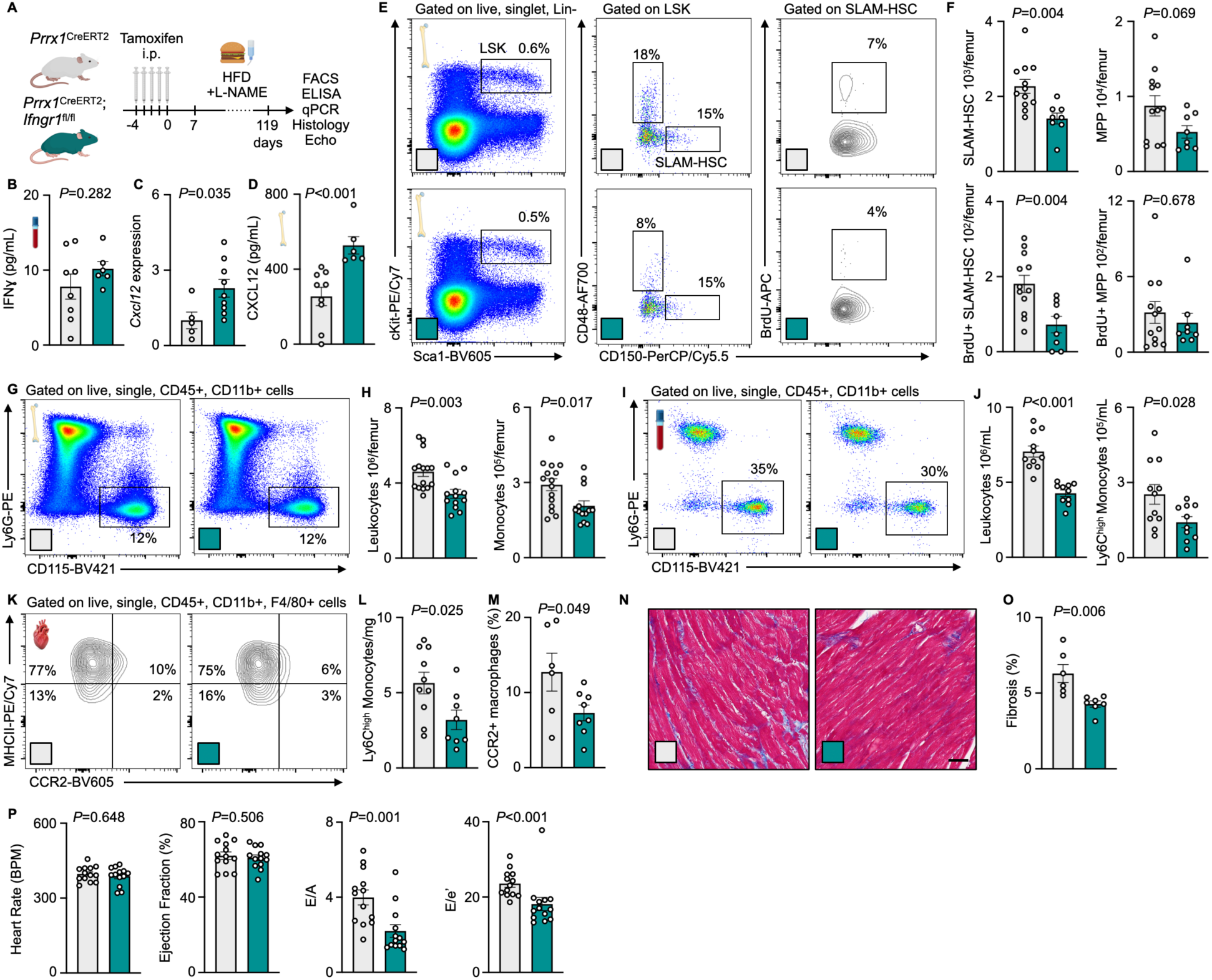
MSC sensing of IFNɣ increases myelopoiesis in HFpEF. (**A**) Experimental outline of *Prrx1*^CreERT2^ vs. *Prrx1*^CreERT2^;*Ifngr1*^fl/fl^ mice with HFpEF. (**B-D**) Protein levels of IFNɣ in the blood, *Cxcl12* gene expression in sorted MSCs and CXCL12 in bone marrow plasma of *Prrx1*^CreERT2^;*Ifngr1*^fl/fl^ mice compared with *Prrx1*^CreERT2^ controls, both with HFpEF (n=5-9 per group, unpaired student’s t-test or two-tailed Mann-Whitney test, depending on normality). (**E**) Gating strategy to identify SLAM-HSC and MPP as well as their proliferation, by BrdU incorporation. (**F**) Quantification of SLAM-HSC and MPP number and proliferation in *Prrx1*^CreERT2^ vs. *Prrx1*^CreERT2^;*Ifngr1*^fl/fl^ mice with HFpEF (n=8-12 per group, unpaired student’s t-test or two-tailed Mann-Whitney test, depending on normality). (**G-H**) Flow cytometry gating and enumeration of bone marrow monocytes in *Prrx1*^CreERT2^;*Ifngr1*^fl/ fl^ mice and *Prrx1*^CreERT2^ controls, both with HFpEF (n=13-15 per group, two-tailed Mann-Whitney test). (**I-J**) Gating and enumeration of blood monocytes in *Prrx1*^CreERT2^;*Ifngr1*^fl/fl^ and *Prrx1*^CreERT2^ controls, both with HFpEF (n=10-11 per group, unpaired student’s t-test or Mann-Whitney test, depending on normality). (**K**) Gating to identify CCR2+ cardiac macrophage populations. (**L**) Ly6C^high^ monocyte quantification in *Prrx1*^CreERT2^;*Ifngr1*^fl/fl^ and *Prrx1*^CreERT2^ controls, both with HFpEF (n=8-10 per group, unpaired student’s t-test). (**M**) Quantification of CCR2+ macrophage percentage in *Prrx1*^CreERT2^;*Ifngr1*^fl/fl^ and *Prrx1*^CreERT2^ controls, both with HFpEF (n=6-8 per group, unpaired student’s t-test). (**N**) Masson’s trichrome staining of heart sections from *Prrx1*^CreERT2^ control mice vs. *Prrx1*^CreERT2^;*Ifngr1*^fl/fl^ mice post-HFpEF. Scale bar = 50 µm. (**O**) Quantification of interstitial fibrosis in *Prrx1*^CreERT2^;*Ifngr1*^fl/fl^ mice and *Prrx1*^CreERT2^ control mice with HFpEF (n=6-7 per group, unpaired student’s t-test). (**P**) Diastolic function measured by Doppler echocardiography in *Prrx1*^CreERT2^;*Ifngr1*^fl/fl^ mice compared with *Prrx1*^CreERT2^ controls, both with HFpEF (n=13 per group, two-tailed Mann-Whitney test).

### Visceral adipose tissue CD8+ T cells are the primary source of IFNɣ in mice with HFpEF

After seeing an MSC-relayed IFNɣ response in HFpEF, we next aimed to determine the source of IFNɣ in mice with HFpEF. We exposed IFNɣ reporter (GREAT) mice to HFpEF and studied eYFP reporter gene expression levels across immune cells in major organs, comparing to healthy control GREAT mice (Figure 6A, Figure S7). Comparing eight organs and various cell types, we found that CD8+ T cells in visceral adipose tissue (VAT) had the most expanded eYFP expression between HFpEF and control mice. VAT from HFpEF mice had increased numbers of both CD8+ T cells and their eYFP+ fraction (Figure 6B and 6C), findings that agree with studies in obese humans.^43^ We then depleted CD8+ T cells in mice with HFpEF to determine if these cells were the dominant source of IFNɣ (Figure 6D). After confirming successful antibody CD8+ T cell depletion (Figure 6E and 6F), we quantified circulating IFNɣ levels by ELISA and found them to be significantly reduced in CD8+ T cell-depleted mice (Figure 6G). CXCL12 levels in the bone marrow of CD8+ T cell-depleted mice rose (Figure 6H). Decreased CD8+ T cells and blood IFNɣ, resulting in elevated bone marrow CXCL12, led to a concomitant drop in circulating monocytes (Figure 6I and 6J).

**Figure 6:**
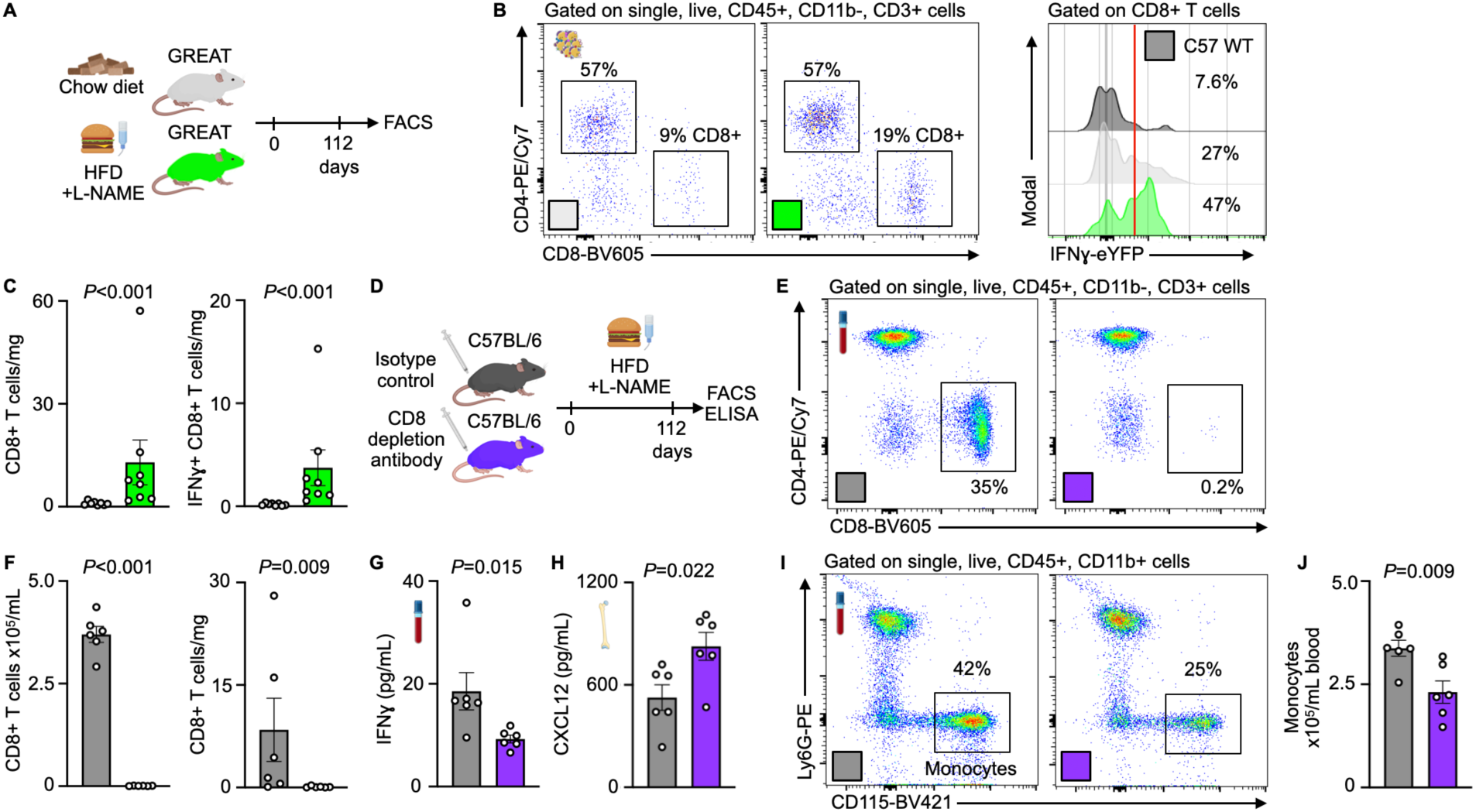
CD8+ T cells in visceral adipose tissue are the IFNɣ source in HFpEF. (**A**) Experimental outline of HFpEF in IFNɣ reporter mice (GREAT). (**B**) Flow cytometry gating for IFNɣ expression in CD8+ T cells from visceral adipose tissue. (**C**) Total and IFNɣ-producing CD8+ T cell numbers in visceral adipose tissue (VAT) in GREAT mice with HFpEF compared to GREAT control mice without HFpEF (n=8 mice per group, two-tailed Mann-Whitney test). (**D**) Experimental outline of CD8+ T cell depletion in mice with HFpEF. (**E**) Flow cytometry gating for CD8+ T cells in blood. (**F**) CD8+ T cell numbers in blood and VAT following CD8+ T cell depletion (n=6 per group, unpaired student’s t-test). (**G**) IFNɣ levels in blood in mice with HFpEF with and without CD8+ T cell depletion (n=6 per group, two-tailed Mann-Whitney test). (**H**) CXCL12 levels in the bone marrow plasma of mice with HFpEF with and without CD8+ T cell depletion (n=6 per group, unpaired student’s t-test). (**I**) Gating for blood monocytes in CD8+ depleted mice vs. isotype control mice. (**J**) Monocyte levels in the blood of CD8+ depleted mice vs. isotype controls (n=6 per group, unpaired student’s t-test).

### MSC-specific IFNɣ response is specific for HFpEF and does not occur in heart failure with reduced ejection fraction (HFrEF)

IFNɣ signaling increases in multiple forms of heart failure and can act on myriad cell types.^44^ To compare the contribution of bone marrow MSC sensing IFNɣ in HFpEF versus HFrEF, we performed scRNA-seq on bone marrow niche cells from mice with myocardial infarction-induced HFrEF and HFpEF (Figure 7A). For this analysis, bone marrow niche cells associated with a major cell type were first grouped into larger clusters (Figure 7B). Scatterplot analysis of MSC GSEA highlighted a unique IFNɣ inflammatory signature in HFpEF versus control MSC as compared to HFrEF versus control MSC (Figure 7C). When directly comparing HFpEF MSC to HFrEF MSC via GSEA, gene sets associated with IFNɣ response and inflammation were significantly enriched in the HFpEF group (Figure 7D and 7E). Top DEGs in MSC highlighted elevated IFNɣ sensing and response in HFpEF compared to HFrEF (Figure 7F). These data identify IFNɣ sensing by MSC as specific to HFpEF when compared with HFrEF.

**Figure 7:**
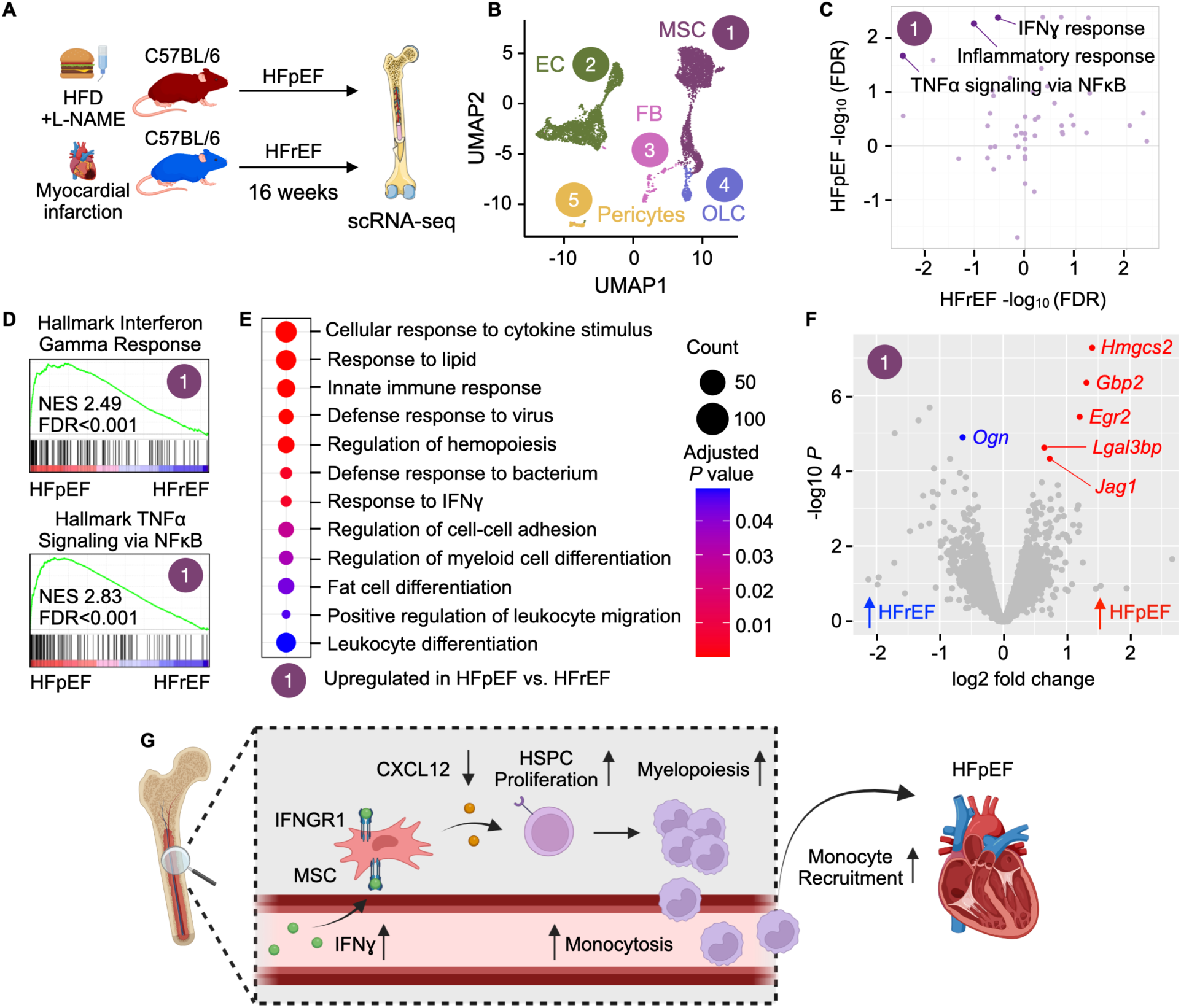
Single cell analysis of bone marrow MSC in HFpEF vs. HFrEF. (**A**) Experimental outline for scRNA-seq of bone marrow stromal cells in control, HFpEF and HFrEF mice. (**B**) UMAP identifying major cell types. (**C**) Scatterplot comparing Hallmark gene sets enriched in MSC in HFpEF vs. control and HFrEF vs. control. A positive −log_10_(FDR) value over 1.3 indicates a statistically significant increase in expression. (**D**) GSEA in MSC in HFpEF vs. HFrEF. (**E**) GOBP gene sets enriched in HFpEF vs. HFrEF in MSC. (**F**) Volcano plot highlighting the top DEGs in MSC in HFpEF vs. HFrEF. (**G**) The bone marrow response in HFpEF: MSC respond to increased IFNɣ levels by lowering levels of CXCL12, thereby influencing HSPC activity and downstream production of pathogenic monocytes.

## Discussion

Obesity is a dominant risk factor for HFpEF, and both obesity and HFpEF are expanding rapidly worldwide. The mechanisms linking obesity to HFpEF are considered a key knowledge gap,^45^ and understanding them may provide a path for identifying much needed therapeutic interventions. Our study describes how adipose tissue, the immune system and the bone marrow interact during the genesis of HFpEF in obese, hypertensive mice: adipose tissue-derived CD8+ T cells signal to the hematopoietic niche via the inflammatory cytokine IFNɣ, which increases systemic and cardiac supply of disease-promoting myeloid cells (Figure 7G). Previously, HFpEF has been shown to involve organs beyond the heart; here we highlight, for the first time, the bone marrow as playing a potential pathogenic role in HFpEF.

HFpEF is thought to be a syndrome with heterogeneous root causes. Thus, our findings may apply to a subset of patients whose risk factor profile is similar to that of mice studied here.^39^ Of note, obesity and hypertension are highly prevalent in HFpEF patients. Nevertheless, we speculate that IFNɣ signaling to hematopoietic niche cells may be most important in (or limited to) obese HFpEF patients, a notion that is also supported by previous work indicating that in obesity, adipose tissue T cells amplify inflammation locally and in the vasculature via IFNɣ signaling.^46^ While we here show the importance of T cell-derived IFNɣ signaling from adipose tissue to the bone marrow in HFpEF, other signaling pathways likely contribute to elevated hematopoiesis in this setting. For instance, adipose tissue macrophages produce IL-1β that is directly sensed by myeloid progenitors in the bone marrow.^11^ In addition, T cells may also directly affect the myocardium in mice with HFpEF.^47, 48^

Comparing the hematopoietic microenvironments in HFpEF and HFrEF, which also associates with increased myelopoiesis,^49^ leukocytosis and expanded inflammatory myocardial leukocytes, indicates that specific, diverging pathways trigger leukocyte oversupply in each condition. This insight aligns with the clinical notion that these forms of heart failure require different therapy, i.e. that efficient HFrEF drugs fail to improve HFpEF.^50^ In our study, mice with HFrEF did not have hypertension and were not obese, which is perhaps a straightforward explanation for the disparate bone marrow transcriptomics results. The hematopoietic retention and quiescence factor CXCL12 is also reduced after myocardial infarction; however, in that context it is thought to result from sympathetic nervous signaling rather than IFNɣ signaling.^51^ Indeed, we found that IFNɣ-related pathways were much more prevalent in mice with HFpEF than in those with HFrEF. This dichotomy between heart failure types may be less pronounced in human patients, as hypertension and obesity are risk factors in both conditions.

We found that deleting the IFNɣ receptor from MSC decreased hematopoiesis not only in mice with HFpEF but also in the steady state, thereby indicating that sensing the inflammatory cytokine IFNɣ by MSC functions as a rheostat for baseline hematopoiesis. Some reports suggest that tonic IFNɣ signaling augments steady-state HSPC activity;^52^ however, others found that direct sensing of IFNɣ by HSPC inhibits hematopoiesis while favoring certain blood cell lineages.^53^ We believe that these seemingly contradictory reports can be reconciled by a model in which direct IFNɣ signaling to HSPC limits cell proliferation while indirect signaling via MSC, relayed via lowered CXCL12, increases HSPC proliferation. We speculate that such a dual system may help orchestrate emergency hematopoiesis during which quickly producing short-lived innate immune cells takes precedence over generating longer-lived red blood cells and lymphocytes. Note that, in the setting of viral infection, CD8 T cells provide IFNɣ to bone marrow stromal cells, which up-regulate IL-6 production and thus expand myelopoiesis.^22^ Whether inhibitory and tonic IFNɣ actions on hematopoiesis occur in separate micro-environments that engender specific blood lineages is an interesting open question.

IFNɣ is a major pro-inflammatory signal after infection and injury, contexts in which it, for instance, activates lymphocytes and primes macrophages, cells that mount defenses against viral, fungal or bacterial challenges.^54^ Binding of IFNɣ to its receptor activates JAK-STAT signaling and expands expression of many interferon-stimulated genes, including those for chemokines and proteins important for phagocytosis, killing infectious agents and antigen presentation. Humans with genetic IFNɣ signaling disruption are prone to various infections.^54^ Given its essential defense functions, broadly neutralizing IFNɣ is likely not a viable strategy in HFpEF. Instead, future drug development may either aim upstream to inhibit metabolic pathways that increase IFNɣ production in CD8 T cells or pursue agonists for the CXCL12 receptor CXCR4 which is expressed by bone marrow HSPC, among other cells. Once translated to the clinical setting, preferential drug delivery to specific bone marrow niches may enable interference with the pathway without compromising defense.^55^

## Resource Availability

Sequencing data were deposited in NCBI’s Gene Expression Omnibus and are accessible through GEO Series accession number GSE292276.

## Acknowledgments

We would like to thank the HSCI-CRM Flow Cytometry Core for assistance with cell sorting, the Histopathology Research Core at MGH for assistance with paraffin embedding and histological staining, BioRender (MGB License) for cartoon components and Kaley Joyes for manuscript editing. We acknowledge Yoshiko Iwamoto for her expert advice on histology. This work was funded by the National Institutes of Health (HL142494, HL176359, HL007208, HL760439, HL15073, HL166538, HL170058), Deutsche Forschungsgemeinschaft (DFG) #453989101 SFB 1525, Mercator fellow M.N, and DFG #530157297 (S.P.).

## Author Contributions

K.M., D.R. and M.N. conceived the study.

K.M., D.R. and M.N. designed the experiments.

K.M., I.L., D.R., A.P., H.S., N.K., N.M., S.P., C.G.M., K.T. and S.U. performed the experiments. K.M., I.L., K.N., M.H. and M.N. analyzed data.

K.M., I.L. and M.N. made figures.

M.S. identified patients for consent.

M.C-K. and A.K. confirmed patient consent and collected human samples.

V.K., R.J., X.M. and J.W. processed human samples.

P.v.G. and A.K. designed the human bone marrow collection and provided de-identified clinical data.

K.M., D.R., A.P., H.S., J.R., K.N., M.H. and M.N. discussed results and strategy.

M.N. directed the study.

K.M. and M.N. wrote the manuscript with input from all authors.

## Methods

### Human bone marrow samples

Human sternal bone marrow were collected from patients undergoing cardiac surgery at Massachusetts General Hospital. Procedures performed at the time of collection included: coronary artery bypass surgery, replacement or repair of the aortic valve, replacement or repair of the tricuspid valve, replacement or repair of the mitral valve, pericardiectomy, replacement of ascending aorta or Maze. A subset of patients underwent multiple procedures simultaneously. Human tissue collection was conducted according to the Declaration of Helsinki and was approved by the Mass General Brigham Institutional Review Board (2020P004103) with written, informed consent before collection. Patients with HFpEF were enrolled in this study based on confirmed HFpEF diagnosis along with identification of signs and symptoms in previous medical history records (n=7). Patients were not included in the study if they met any of the following exclusion criteria: LVEF<50%, congenital heart disease, severe mitral valve regurgitation, diabetes, acute myocardial infarction or previous heart transplant. Patients were considered as controls if they underwent a cardiac procedure, did not have HFpEF and did not fit the exclusion criteria (n=7).

Bone marrow was collected from patients via sternotomy during open heart surgery. With a spinal fusion curette (size 5), the exposed interior of the sternum was scraped to collect ∼2 mL of bone marrow into a sterile specimen container. To prevent coagulation, 1 mL of heparin (1,000 units/mL, H3149, Sigma) was added to bone marrow samples. To isolate bone marrow plasma, heparinized bone marrow samples were diluted 1:1 with PBS, layered over Lymphoprep^TM^ (18061, Stemcell Technologies) and centrifuged at 1200g for 10 minutes with brake off. The upper plasma layer was then flash frozen and stored for future analyses.

### Mice

Wildtype C57BL/6 (stock 000664), C.129S4(B6)-*Ifng*^tm3.1Lky^/J (GREAT, stock 017580), B6.Cg-Tg(*Prrx1*-cre/ERT2,-EGFP)1Smkm/J (*Prrx1*^CreERT2^, stock 029211), C57BL/6N-*Ifngr1^tm1.1Rds^*/J (*Ifngr1*^fl/fl^, stock 025394), B6.129S4-*Ccr2^tm1lfc^*/J (*Ccr2*^-/-^, stock 004999), B6.129P2 (Cg)- *Cx3cr1^tm^*^2^*^.1(cre/ERT^*^2^*^)Litt^/*WganJ *(Cx3cr1*^CreER^, stock 021160) and B6.Cg-*Gt(ROSA)26Sor^tm9(CAG-tdTomato)Hze^*/J(*Ai9*^fl/fl^, stock 007909) were purchased from the Jackson Laboratory. Genetic crossing and genotyping were performed as described on the Jackson Laboratory website. All experiments were performed with 8- to 12-week old male mice unless otherwise noted. Control groups were age- and sex-matched. Data shown were acquired in at least 2 independent experiments. Mice were housed according to standard group housing guidelines and had unrestricted access to food and water. All animal experiments were approved by the Institutional Animal Care and Use Committee at the Massachusetts General Hospital (2014N000078).

#### Two-hit model of HFpEF

To induce heart failure with preserved ejection fraction as previously described,^29^ mice were fed a high-fat diet (D12492, Research Diets Inc.) and given hypertension-inducing L-NAME hydrochloride (500 mg/L, N5751, MilliporeSigma) in their drinking water for the indicated periods of time.

#### Permanent LAD ligation for heart failure with reduced ejection fraction (HFrEF)

Left anterior descending coronary artery ligation was performed to induce myocardial infarction. Mice were anesthetized with 2% isoflurane, intubated and placed on a ventilator for controlled anesthesia and respiration during surgery. Mice were then placed on a heated surgical pad and depilatory cream was applied to the chest for hair removal. A left thoracotomy was performed in the third or fourth intercostal space to visualize the heart at the level of the left ventricle. A monofilament nylon 8-0 suture was used to permanently ligate the left anterior descending coronary artery, and the thoracic space was then closed with a monofilament nylon 5-0 suture. Analgesia was administered just before surgery and for 3 days after to ensure proper pain management. Mice were then housed for the indicated time periods to allow heart failure to develop.

#### Antibody-based depletion/neutralization

To deplete CD8+ T cells, mice were intraperitoneally (i.p.) injected with 200 µg of *InVivo*MAb anti-mouse CD8a (clone 2.43, BioXCell) or *InVivo*MAb rat IgG2b isotype control, anti-keyhole limpet hemocyanin (clone LTF-2, BioXCell), every 3 days throughout the course of the HFpEF protocol.

#### Tamoxifen injections

Tamoxifen (T5648, MilliporeSigma) was resuspended in corn oil (C8267, MilliporeSigma) at a concentration of 20 mg/mL. Animals received daily doses of 2 mg tamoxifen i.p. over the course of 5 days.

#### Echocardiography

Echocardiography was performed with a Vivo 3100 Imaging System (FUJIFILM VisualSonics) using an MX250s transducer. Mice were anesthetized with 2% isoflurane and heart rates were maintained in a consistent range in order to measure diastolic and systolic function.

#### Tail cuff blood pressure measurements

Blood pressure in conscious mice was measured using a non-invasive tail-cuff system (Kent Scientific) per the manufacturer’s directions.

### Histology

#### Mouse heart processing and analysis

Mice were euthanized via isoflurane inhalation and perfused with 10 mL of cold-PBS through the left ventricle. Hearts were subsequently removed for histological processing. Perfused hearts were fixed in 10% neutral buffered formalin overnight at room temperature, transferred to 70% ethanol for short term storage and then paraffin embedded. Paraffin blocks were sectioned at 5 µm and stained with Masson’s trichrome for downstream fibrosis quantification. Trichrome-stained sections were imaged with a NanoZoomer 2.0RS (Hamamatsu) and analyzed with Fiji. To measure interstitial fibrosis, three fields of view excluding perivascular fibrosis from the left ventricle per mouse were exported at 10X magnification in Nanozoomer NDP.view2 software. The percentage area of interstitial collagen (blue) vs. all tissue was used to calculate interstitial fibrosis.

### Cells

#### Organ and tissue processing

Prior to organ harvest, mice were perfused through the apex of the left ventricle with 10 mL of ice-cold PBS. Organs were then excised, cleaned and micro dissected in preparation for digestion. Micro dissected tissues from heart, lung, liver, spleen, thymus and visceral adipose tissue were digested with 450 U/mL of collagenase I, 125 U/mL of collagenase XI, 60 U/mL of DNase I and 60 U/mL of hyaluronidase (MilliporeSigma) for 60 minutes in a 37C° shaking block. Digested tissue was then triturated and filtered through a 40-µm cell strainer, washed and centrifuged at 340g for 7 minutes to isolate single cells.

To collect peripheral blood, mice were anesthetized with isoflurane and blood was collected via retro-orbital bleeding using heparinized capillary tubes. Red blood cells were lysed using a 1X RBC lysis buffer (BioLegend). Lysed blood was then washed with PBS and spun at 340g for 7 minutes to isolate circulating cells.

To detect leukocytes or HSPCs in mouse bone marrow, both femurs were harvested, cleaned and flushed with PBS. Flushed bone marrow cells were centrifuged at 340g for 7 minutes and resuspended in PBS prior to downstream assays.

To isolate bone marrow niche cells, femurs, tibiae, humeri and pelvic bones were harvested and pooled from two mice for each biological replicate. Bones were flushed with PBS, crushed with a mortar and pestle and enzymatically digested (Dispase II (D4693, MilliporeSigma), DNase I (D5319, MilliporeSigma), Collagenase Type I (07416, StemCell Technologies). Digested bone marrow samples were filtered, washed and centrifuged at 340g for 7 minutes to create a single cell suspension. RBC lysis was then performed and CD45+ cells were subsequently depleted using CD45+ MicroBeads (Miltenyi Biotec) to reduce contamination of RBCs and leukocytes, respectively. Following CD45 depletion, the flow-through of CD45-cells was collected, centrifuged at 340g for 7 minutes and resuspended in PBS for downstream assays.

To label proliferating cells, 100 µL of BrdU (10 mg/mL, 552598, BD Biosciences) was i.p. injected into mice 24 hours prior to harvest.

#### Flow cytometry

All antibody staining steps occurred in PBS on ice. Blood and bone marrow leukocytes were stained with anti-CD45-BV711 (clone 30-F11, BioLegend), -CD11b-BV785 (M1/70, BioLegend), -CD19-APC/Cy7 (6D5, BioLegend), -B220-APC/Cy7 (RA3-6B2, BioLegend), -NK1.1-APC/Cy7 (PK136, BioLegend), -CD3-PerCP/Cy5.5 (17A2, BioLegend), -CD4-PE/Cy7 (GK1.5, BioLegend), -CD8-BV605 (53-6.7, BioLegend), -CD115-BV421 (AFS98, BioLegend), -Ly6G-PE (1A8, BioLegend), -Ly6C-APC (HK1.4, BioLegend) and Live/Dead Fixable Aqua (Thermo Fisher).

In most cases, heart leukocytes were first stained with lineage markers, including anti-B220-PE (RA3-6B2, BioLegend), -CD49b-PE (HMa2, BioLegend), -CD90.2-PE (53-2.1, BioLegend), -Ly6G-PE (1A8, BioLegend), -NK1.1-PE (PK136, BioLegend), -TER119-PE (TER-119, BioLegend) and -CD103-PE (2E7, BioLegend). Cells were then washed and stained with anti-CD45-BV711 (30-F11, BioLegend), -CD11b-APC/Cy7 (M1/70, BioLegend), -F4/80-PE/Cy7 (BM8, BioLegend), -Ly6C-FITC (HK1.4, BioLegend) and DAPI. To identify CCR2+ recruited heart macrophages, we used anti-CD45-BV711 (30-F11, BioLegend), -CD11b-BV785 (M1/70, BioLegend), -CCR2-BV605 (SA203G11, BioLegend), -F4/80-PE/Cy7 (BM8, BioLegend), -Ly6C- APC (HK1.4, BioLegend), -MHCII-APC/Cy7 (M5/114.15.2, BioLegend) and DAPI. To identify a more “resident” cardiac macrophage population, we stained with anti-CD45-BV711 (30-F11, BioLegend), -CD11b-BV785 (M1/70, BioLegend), -CCR2-BV605 (SA203G11, BioLegend), -F4/80-PE/Cy7 (BM8, BioLegend), -Ly6C-APC (HK1.4, BioLegend), -MHCII-APC/Cy7 (M5/114.15.2, BioLegend), -LYVE1-PE (ALY7, Thermo Fisher), -TIMD4-APC (RMT4-54, BioLegend) and Live/Dead Fixable Aqua (Thermo Fisher). For flow cytometry of cardiac fibroblasts, cells were stained with anti-CD45-BV711 (30-F11, BioLegend), -CD31-PerCP/Cy5.5 (390, BioLegend), -Anti-Feeder Cells-APC (mEF-SK4, Miltenyi Biotech), -CD146-PE (ME-9F1, BioLegend), -IFNGR1-BV605 (CD119, GR20, BD Biosciences) and DAPI.

To identify bone marrow HSPCs, bone marrow cells were first stained with lineage markers, including anti-CD3-Biotin (17A2, BioLegend), -CD4-Biotin (GK1.5, BioLegend), -CD8-Biotin (53-6.7, BioLegend), -CD90-Biotin (53-2.1, BioLegend), -CD19-Biotin (6D5, BioLegend), -B220- Biotin (RA3-6B2, BioLegend), -NK1.1-Biotin (PK136, BioLegend), -TER119-Biotin (TER-119, BioLegend), -CD11b-Biotin (M1/70, BioLegend), CD11c-Biotin (N418, BioLegend) and -Gr-1- Biotin (RB6-8C5, BioLegend). Bone marrow cells were then stained with Sca1-BV605 (D7, BioLegend), c-Kit-PE/Cy7 (2B8, BioLegend), CD34-FITC (RAM34, BioLegend), CD16/32-BV711 (93, BioLegend), CD150-PerCP/Cy5.5 (TC15-12F12.2, BioLegend), CD48-AF700 (HM48-1, BioLegend), streptavidin-APC/Cy7 (BioLegend) and Live/Dead Fixable Aqua (Thermo Fisher). To identify bone marrow endothelial and mesenchymal stromal cells, digested bone marrow was first stained with LepR-Biotin (Cat# BAF497, R&D Systems) and then washed and stained with TER119-APC (TER-119, BioLegend), CD45-BV711 (30-F11, BioLegend), Gr-1-PE (RB6-8C5, BioLegend), CD11b-PE (M1/70, BioLegend), CD3-PE (17A2, BioLegend), CD19-PE/Cy7 (6D5, BioLegend), B220-PE/Cy7 (RA3-6B2, BioLegend), CD71-PE/Cy7 (RI7217, BioLegend), CD31- BV605 (390, BioLegend), Sca1-FITC (D7, BioLegend), streptavidin-APC/Cy7 (BioLegend) and DAPI.

To identify proliferating cells by flow cytometry, the APC BrdU Flow Kit (552598, BD Biosciences) was used according to the manufacturer’s protocol.

#### Colony-forming unit assay

Colony-forming unit (CFU) assays were performed according to the manufacturer’s protocol (STEMCELL Technologies). For blood CFUs, 50 uL of RBC-lysed blood was mixed into prepared MethoCult tubes (M3434, STEMCELL Technologies) and subsequently plated into 6- well non-tissue-culture-treated plates. For bone marrow CFUs, 1.5 x 10^5^ bone marrow cells were mixed into prepared MethoCult tubes (M3434) and plated in the same way as above. All samples were incubated at 37C° for 7 days prior to CFU counting.

### ScRNA-seq

#### Tissue preparation

The bones from 2 mice were pooled for each sequencing replicate. Following bone marrow niche cell isolation, cell suspensions were stained with DAPI and Calcein (C3100MP, Thermo Fisher) to identify live, metabolically active cells, as well as those with hematopoietic lineage markers (CD45, TER119, CD19, B220, CD71, Gr-1, CD11b and CD3) to exclude hematopoietic cells. Live (DAPI-), active (Calcein+), lineage^-^ single cells were FACS-sorted and immediately processed with the 10X Genomics Chromium Single Cell 3’ Reagent kit according to the v3 user guide. Generated library quality was validated using an Agilent High Sensitivity DNA chip and sequenced on an Illumina NextSeq 500 system.

#### Data processing

For each sample, the raw sequencing reads were processed with 10x Genomics Cell Ranger 6.1.1, using mm10 as the reference genome, to obtain the count matrix with the number of unique molecular identifiers (nUMI) for each gene in each cell. Within a sample, a cell was considered low-quality and removed if either its nUMI or its number of detected genes (nGene) was lower than the respective sample median by more than three times the median absolute deviation (MAD), or if its percentage of mitochondrial genes contributing to total UMI count (percent.mt) was higher than the sample median by more than three times the MAD. Genes detected in fewer than 10 cells were excluded. Seurat 4.1.0 was used for further analyses,^56–58^ including normalization with v2 version of sctransform,^59^ integration,^57^ principal component analysis and clustering with the top 40 principal components. The leukocyte subset was removed after the initial clustering, and the remaining cells were reclustered following the same normalization and integration workflow described above.

#### Differential gene expression and gene set enrichment analyses

Pseudobulk differential expression analysis^60^ was carried out with a quasi-likelihood F-test based on the generalized linear model implementation of edgeR 3.34.1^61–63^ for each cluster to detect differentially expressed genes between cells in different conditions.

The GSEAPreranked tool from GSEA 4.2.2^64, 65^ was used to perform gene set enrichment analysis of gene sets in the molecular signatures database MSigdB 7.5.1.^64, 66^ The gseGO function of R package clusterProfiler 4.1.4^67^ was used to create the dot plots in Figures 3F and 7E.

### Molecular Biology

#### RNA isolation and qPCR

RNA was isolated using the RNeasy Mini Kit (74104, Qiagen) or the Plus Micro Kit (74034, Qiagen), depending on cellular numbers. For reverse transcription, the High-Capacity RNA-to-cDNA Kit (4387406, Applied Biosystems) was used. Quantitative qPCR was performed using TaqMan gene expression assays with TaqMan Fast Universal PCR Master Mix (4366072, Applied Biosystems) and specific primers for *Cxcl12* (Mm00445553_m1, FAM-MGB) and *Gapdh* (Mm99999915_g1, VIC-MGB, Thermo Fisher). Data were acquired on a CFX Opus 96 Real-Time PCR System (Bio-Rad). Samples were run in duplicate and gene expression was normalized to *Gapdh*.

#### ELISA

##### Mouse

IFNɣ and CXCL12 were measured in blood serum and bone marrow plasma by using the Mouse IFN-gamma DuoSet ELISA (DY485-05, R&D Systems) and Mouse SDF1/CXCL12 ELISA Kit (ab100741, Abcam), respectively. To isolate serum, 500 uL of whole blood was collected via cardiac puncture, stored at room temperature for 20 minutes to facilitate clot formation and then centrifuged at 5,000g for 5 minutes. The serum supernatant was collected and stored at −80C until analysis. To isolate bone marrow plasma, both tibiae were harvested and extensively cleaned to remove overlying muscle and traces of liquid on the bone surface. Tibiae were then cut through the diaphysis and centrifuged at 5,000g for 5 minutes. The liquid fraction (bone marrow plasma) was then separated from the cell pellet and stored at −80C until analysis.

##### Human

CXCL12 was measured in bone marrow plasma from patient sternal bone marrow samples using the Human CXCL12/SDF-1 alpha ELISA kit (DSA00, R&D Systems) according to the manufacturer’s protocol. Prior to analysis, samples were spun at 10,000g for 10 minutes to remove any platelet contamination.

### Statistical analysis

Results are expressed as mean ± SEM. Statistical analyses were performed using GraphPad Prism 10. Normality was assessed using the Shapiro-Wilk normality test. For analyses between two groups, either the parametric unpaired two-tailed student’s t-test or non-parametric two-tailed Mann-Whitney test was used, depending on normality.

---

**Figure S1:**
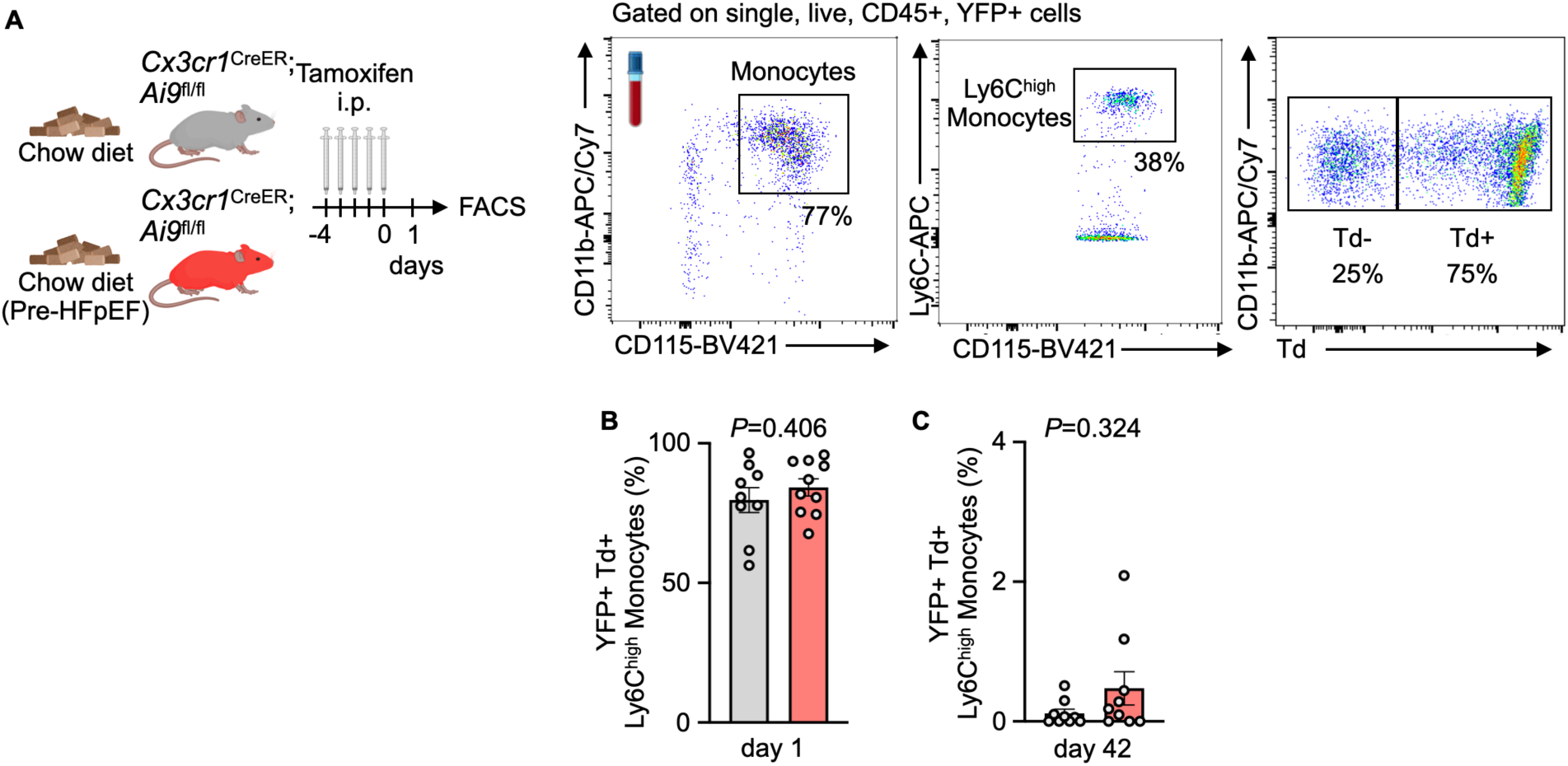
Baseline characterization of *Cx3cr1*^CreER^;*Ai9*^fl/fl^ mice, related to Figure 2. (**A**) Experimental outline and gating strategy of baseline fate mapping experiment in *Cx3cr1*^CreER^;*Ai9*^fl/fl^ mice. (**B-C**) Quantification of baseline and pre-HFpEF YFP+ Td+ Ly6C^high^ monocyte frequency in *Cx3cr1*^CreER^;*Ai9*^fl/fl^ mice in the blood (n=9-10 per group, unpaired student’s t test or two-tailed Mann-Whitney test depending on normality).

**Figure S2:**
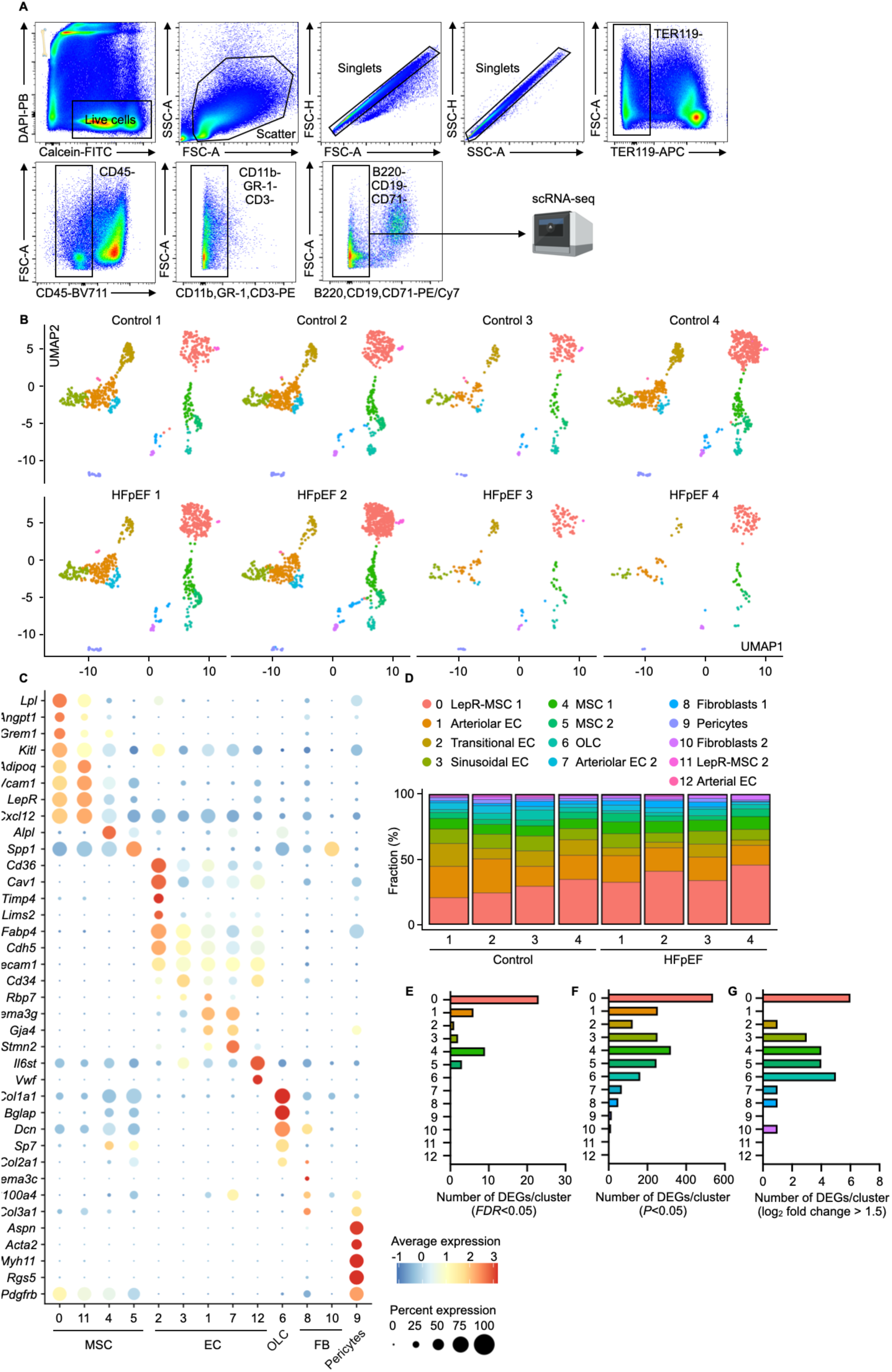
Identification and characterization of scRNA-seq clusters, related to Figure 3. (**A**) Gating strategy for sorting bone marrow niche cells for scRNA-seq. (**B**) UMAPs of individual samples from control and HFpEF experimental groups. (**C**) Cluster defining markers used to determine cellular identity. In each dot, color denotes average gene expression(red: high, blue: low) and size indicates the percentage of cells which express the gene. (**D**) Fraction of cells belonging to each cluster across individual samples. (**E-G**) The number of differentially expressed genes (DEGs) in HFpEF mice vs. controls, split by cluster, shown with unadjusted *P*-values<0.05 or log_2_ fold change > 1.5.

**Figure S3:**
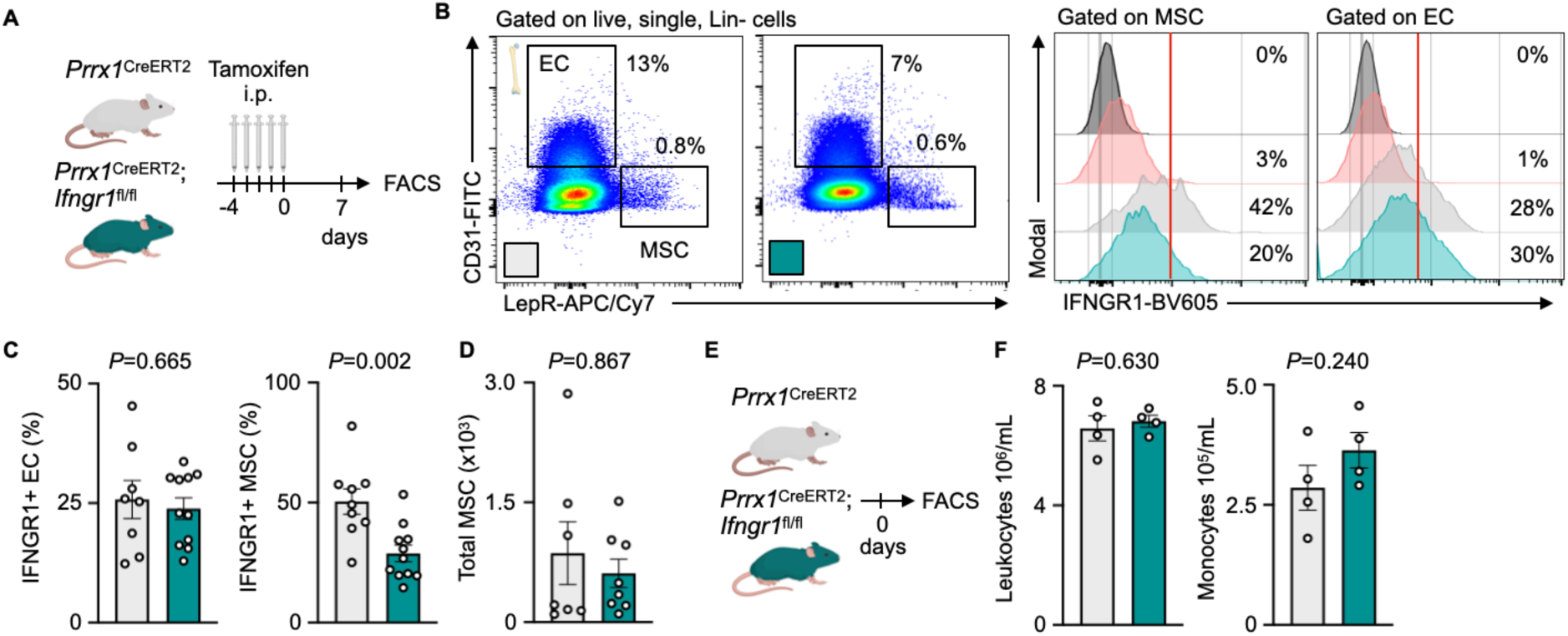
Baseline characterization of *Prrx1*^CreERT2^;*Ifngr1*^fl/fl^ mice, related to Figure 4. (**A**) Experimental outline to measure IFNGR1 expression by flow cytometry. (**B**) Gating strategy to identify IFNGR1+ bone marrow LepR+ MSC and endothelial cells (EC). Black histogram = unstained control, red histogram = IgG2a isotype control. (**C**) IFNGR1 surface expression in bone marrow LepR+ MSC and EC in *Prrx1*^CreERT2^;*Ifngr1*^fl/fl^ vs. *Prrx1*^CreERT2^ mice (n=8-12 per group, unpaired student’s t-test). (**D**) Total number of LepR+ MSC in *Prrx1*^CreERT2^;*Ifngr1*^fl/fl^ vs. *Prrx1*^CreERT2^ mice (n=7-8 per group, two-tailed Mann-Whitney test). (**E**) Experimental outline to measure leukocyte levels prior to tamoxifen administration. (**F**) Leukocyte and monocyte levels in *Prrx1*^CreERT2^;*Ifngr1*^fl/fl^ vs. *Prrx1*^CreERT2^ mice prior to tamoxifen administration (n=4 per group, unpaired student’s t-test).

**Figure S4:**
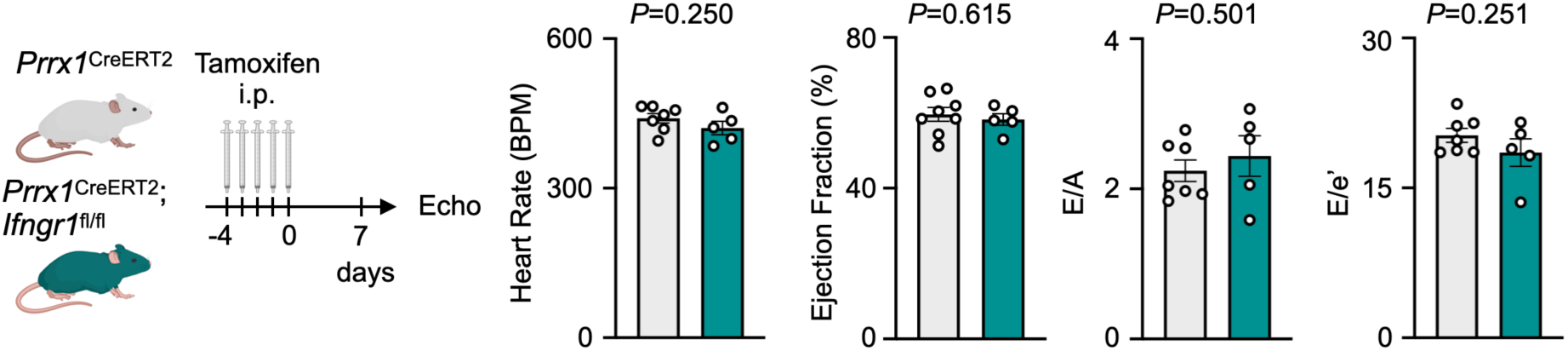
Baseline cardiac function of *Prrx1*^CreERT2^;*Ifngr1*^fl/fl^ mice, related to Figure 4. Systolic and diastolic cardiac function in *Prrx1*^CreERT2^;*Ifngr1*^fl/fl^ vs. *Prrx1*^CreERT2^ mice as measured by Doppler echocardiography (n=5-7 per group, unpaired student’s t-test).

**Figure S5:**
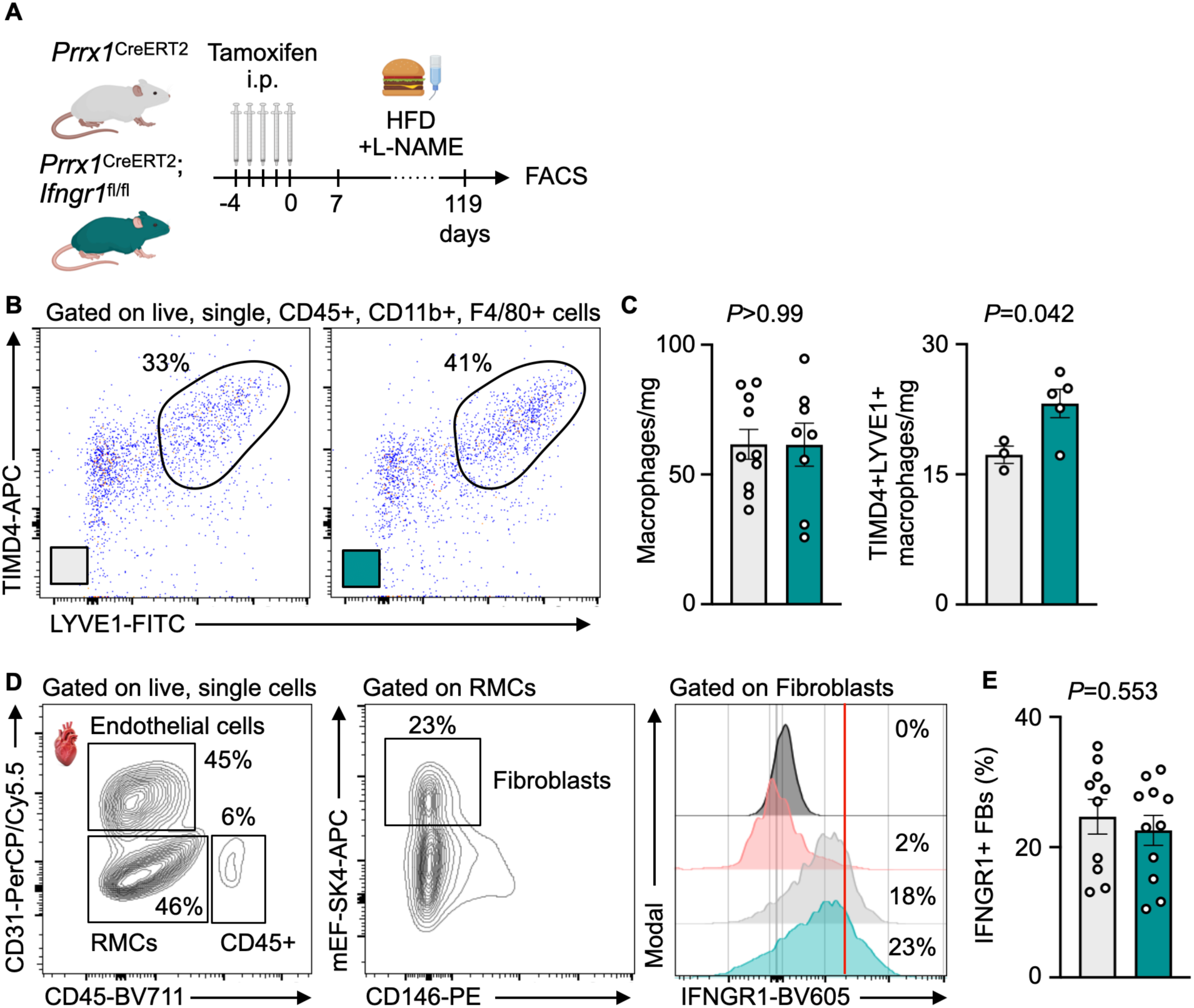
Cardiac macrophage phenotype shift in *Prrx1*^CreERT2^;*Ifngr1*^fl/fl^ mice post-HFpEF, related to Figure 5. (**A**) Experimental outline to characterize cardiac macrophage and fibroblast populations in *Prrx1*^CreERT2^;*Ifngr1*^fl/fl^ vs. *Prrx1*^CreERT2^ mice post-HFpEF. (**B**) Gating strategy to identify TIMD4+LYVE1+ macrophages. (**C**) Total macrophage (n=8-10 per group, unpaired student’s t-test) and TIMD4+LYVE1+ macrophage abundance quantification (n=3-5 per group, unpaired student’s t-test). (**D**) Gating strategy to identify IFNGR1+ cardiac fibroblasts. Black histogram = unstained control, red histogram = IgG2a isotype control. RMCs = Resident mesenchymal cells. (**E**) IFNGR1 surface expression in cardiac fibroblasts in *Prrx1*^CreERT2^;*Ifngr1*^fl/fl^ vs. *Prrx1*^CreERT2^ mice (n=7-8 per group, unpaired student’s t-test).

**Figure S6:**
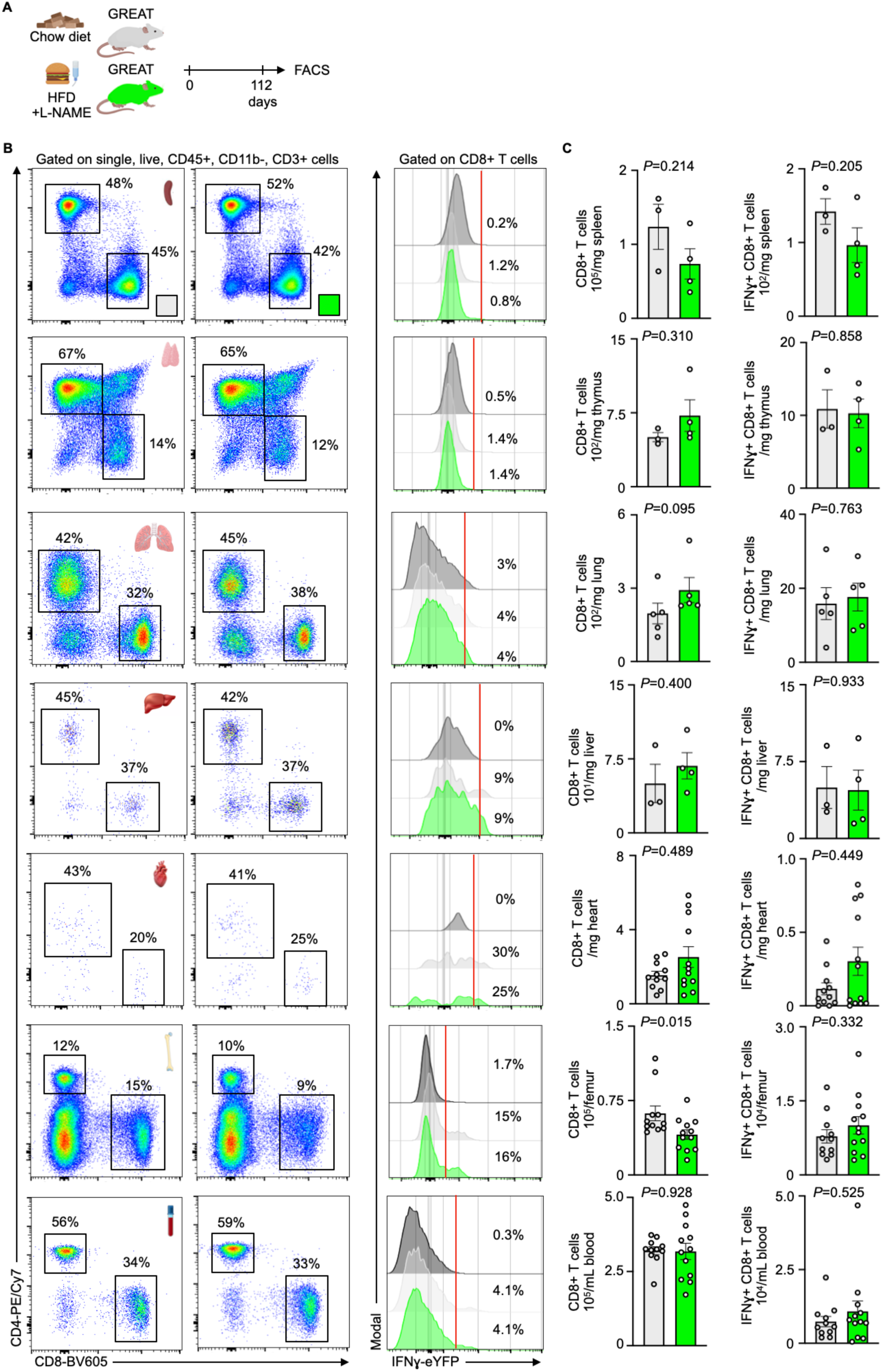
Distribution of IFNɣ+ CD8+ T cells in HFpEF, related to Figure 6. (**A**) Experimental outline of HFpEF in IFNɣ reporter mice (GREAT). (**B**) Gating strategies to identify CD8+ T cells and IFNɣ expression levels in spleen, thymus, lung, liver, heart, bone marrow and blood. (**C**) Quantification of total and IFNɣ+ CD8+ T cell numbers from spleen (n=3-4 per group), thymus (n=3-4 per group), lung (n=5 per group), liver (3-4 per group), heart (n=11-12 per group), bone marrow (n=11-12 per group) and blood (n=11-12 per group), respectively (unpaired student’s t-test or two-tailed Mann-Whitney test, depending on normality).

**Table S1:**
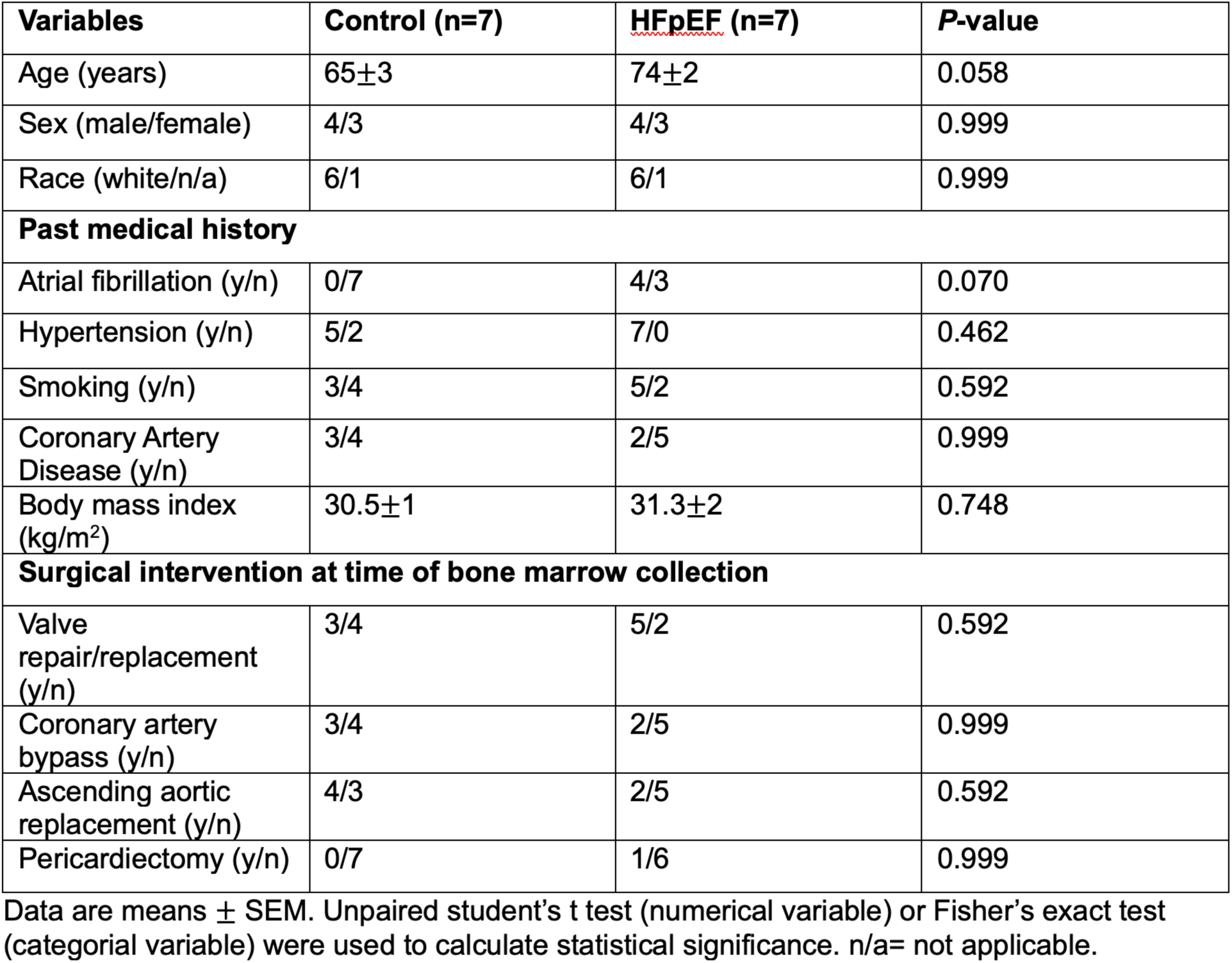
Patient demographics and clinical information.

